# Accordion-like collagen fibrils suggested by P-SHG image modeling : implication in liver fibrosis

**DOI:** 10.1101/420265

**Authors:** D. Rouède, E. Schaub, J-J. Bellanger, F. Ezan, F. Tiaho

## Abstract

Second-order non-linear optical anisotropy parameter *ρ* = *χ*_33_ / *χ*_31_ is calculated for collagen-richt issues considering both a single dominant molecular hyperpolarizability tensor element *β*_333_ = *β* at single helix level and a priori known submicrometric triple helical organization of collagen molecules. Modeling is further improved by taking account of Poisson photonic shot noise of the detection system and simple supra-molecular fibrillar arrangements in order to accurately simulate the dispersion of *ρ* values in collagen-rich tissues such as tendon, skin and liver vessels. From combined P-SHG experiments and modeling, we next correlate experimental and theoretical statistical distributions of *ρ*. Our results highlight that the dispersion of experimental *ρ* values is mainly due to (i) Poisson photonic shot noise in tendon and skin, which proves to have a preponderant effect in P-SHG experiments (ii) variance of supercoil angles of accordion-like fibrils in vessels that is further reduced during the development of liver fibrosis therefore contributing to the rigidity of the tissue. These results open new avenue for future modeling correlating the dispersion of *ρ* values in P-SHG experiments and the fibrillar architecture as well as the mechanical stiffness of patho-physiological extracellular matrices in collagen tissues.

## INTRODUCTION

Second harmonic generation (SHG) microscopy is a label-free technique that relies on a nonlinear optical interaction with hyperpolarizable non-centrosymmetric endogenous fibrillar proteins like collagen and myosin causing scattered coherent radiation at twice the fundamental frequency (1-4). Polarization dependence of SHG (P-SHG) microscopy is gaining increase popularity for investigating fibrillar collagen-rich tissues with the desire to extract as much structural information in physiological as well as in disease state. Organization of collagen tissue using P-SHG is usually described by anisotropy parameter coefficient *ρ* = *χ*_33_ / *χ*_31_ that is defined by the ratio of the two independent second-order nonlinear optical susceptibility tensor coefficients *χ* _33_ and *χ*_31_ that are involved in the nonlinear optical interaction (5-13).

We previously proposed a simple reliable and fast linear least square (LLS) fitting method to process P-SHG images at pixel-resolution (14). More recently, we extended this method to retrieve the pixel-resolved sub-microscopic hierarchical organization (helical, triple-helical) of nonlinear molecules by correlating the experimental and theoretical statistical distributions of *ρ* values through a Monte Carlo simulation taking into account the background Poisson photonic shot noise of the detectors (15). However, we failed to explain rigorously the dispersion of *ρ* values in some tissues such as mouse liver vessels.

The aim of the present article is to solve this problem. For this purpose, we first model the distribution of *ρ* values considering Poisson photonic shot noise of the detectors and different fibrillar architectures (intra and inter pixel arrangements of straight or supercoil fibrils). Then by correlating experimental and theoretical results, we find that the dispersion of *ρ* values is dominated by Poisson photonic shot noise in collagen tendon and skin characterized by quasi-ordered fibrils. However, in liver vessels characterized by supercoiled fibrils, we find that the dispersion of *ρ* values is dominated, in addition to shot noise effect, by pixel-to-pixel variance of supercoil angles suggesting the formation of accordion-like supercoiled fibrils. Moreover, development of fibrosis results in reduction of this inter pixel dispersion therefore explaining the increase rigidity of the vessels.

## MATERIALS AND METHODS

### Preparation of biological samples

Samples from physiological and pathological tissues were prepared as previously reported and summarized as follows (15). Physiological samples were performed with rat and mouse collagen-rich tissues. Adult Wistar rats (200-300g) were euthanized by CO_2_ inhalation. Tissues were dissected, fixed over night with 4% paraformaldehyde in phosphate buffer saline at 4°C, and rinsed at least three times with PBS. For SHG imaging, dissected pieces (100-200 μm thickness) of tissues were mounted in 50% glycerol-PBS solution and stabilized between two coverslips sealed with nail polish. For pathological samples, wild-type C57 black 6 mice received intraperitoneal injection of either 0,5 μg CCl_4_ (Sigma, St Louis, MO) dissolved in oil in order to induce liver fibrosis or the olive oil solvent for control mice. Injections were performed 3 times (D1, D3 and D7) the first week followed by single injection every week lasting 8 weeks (W1-W8). Mice were euthanized at D1 or at W10, livers were harvested and fixed in 4% paraformaldehyde. For SHG-imaging, 10 μm thick sections of liver slices mounted between microscope glass slides and coverslips were obtained from the Biosit-H2P2 core (histopathologie.univ-rennes1.fr/) according to established standard procedure. All rats and mice were cared for in accordance to the “Guide for the Care and Use of Laboratory Animals” (Directive 2010/63/UE).

### Image acquisition and analysis

Images were acquired on a custom made SHG microscope part of the BIOSIT facility https://biosit.univ-rennes1.fr. It is based on an inverted microscope (IX71, Olympus, Japan) and the laser source is a tunable IR 80MHz femtosecond Ti:Sa laser (MAITAI, Spectra Physics). Polarization of the beam is controlled by two achromatic half and quarter wave plates located upstream of the scanning mirrors, and mounted on motorized rotary stages (PR50CC, Newport). High NA water immersion objective (Olympus UPLSAPO 60XW NA=1.20, WD = 0.28 mm) was used for applying 10-20 mW of 740 nm excitation at the sample. PSF was obtained from 0.17 μm diameter fluorescent micro beads (Molecular Probes PS-Speck Microscope Point Source Kit (P7220)) and estimated to be 0.4×1.2 μm. SHG signal was collected in forward direction using high NA objective (Olympus, LUMFl 60XW, NA = 1.1). SHG signal is detected using high sensitivity single photon GasAsP photomultipliers (H7421-40, Hamamatsu). Photons are counted by a general purpose acquisition USB module (NI-USB 6363, National Instruments). It is important to note that in addition to tissue clearing obtained with 50% glycerol treatment, all polarization dependent SHG images were acquired at a depth < 20 μm from the tissue surface to minimize depolarization (16) and birefringence effects (17, 18). All experimental P-SHG stacks were obtained for input polarization angles uniformly distributed between 0° and 180° with 20° increments. This incremental step is a good sampling compromise for accurate acquisition of P-SHG parameters and quality of the recording, avoiding motion artifact. Image analysis and simulations were performed using MALAB (MathWorks, Natick, MA, USA). Total image processing time for a 512 × 512 pixels image including LLS fitting and Kolmogorov-Smirnov test procedure is about 30 s.

## THEORY

Second harmonic electric fields ***E***^*2ω*^ emitted at 2ω originate from nonlinear polarization ***P***^*2ω*^= *χ*^(2)^***E***^*ω*^ ***E***^*ω*^induced by mixing of intense electric fields ***E***^*ω*^at ω in a medium characterized by a macroscopic second-order nonlinear optical susceptibility *χ* ^(2)^. Our strategy to calculate *χ* for each pixel is first to calculate the second order nonlinear optical susceptibility 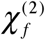 of an individual fibril and then to add the contributions of all the fibrils inside the pixel. 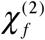 can be obtained from the associated molecular hyperpolarizability tensor by averaging the *β*^(2)^ contributions of all nonlinear dipoles of the fibril. Throughout the rest of the document, we assume that *β*^(2)^ has a single coefficient *β* that has been verified experimentally on 12 different tissues (see Supporting Material S1) and that *β* is dominated by the peptide bonds along the collagen single helix scaffold.

Mechanism of fibril formation is well known (19), it is a hierarchical helical organization process of peptide bonds related *β* as schematized in Fig. 1. Starting from *β*(Fig. 1 a), the hierarchical organization is a multi-step process. The first process is the formation of a single helix (H) polypeptide chain around single helix axis z_1_ characterized by a polar angle *θH* (Fig. 1 b). The second process is the formation of a triple helix (3H) around triple helix axis z_2_ characterized by a polar angle *θ*_3*H*_ (Fig. 1 c) and resulting from the association of three polypeptide chains called tropocollagen (collagen molecule). Staggered and cross-linked assembly of these collagen molecules explain the formation of straight fibrils as in tendon (19). A supercoil (SC) process (Fig. 1 d) is taken into account in dermis where triple helical tropocollagen is twisted at constant polar angle *θ*_*SC*_around supercoiled fibril axis z_3_, forming supercoiled micro fibrils or fibrils (19). Another process may be considered if the fibril is tilted (T) with a polar angle *θ*_*T*_ relative to the microscope stage (Fig. 1 e and f). Taking account of the hierarchical four processes, we found after calculation (see Supporting Material S2) that only three tensor elements 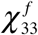 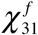 and 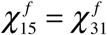 of 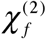 mainly contribute to the SHG signal, and they are given by

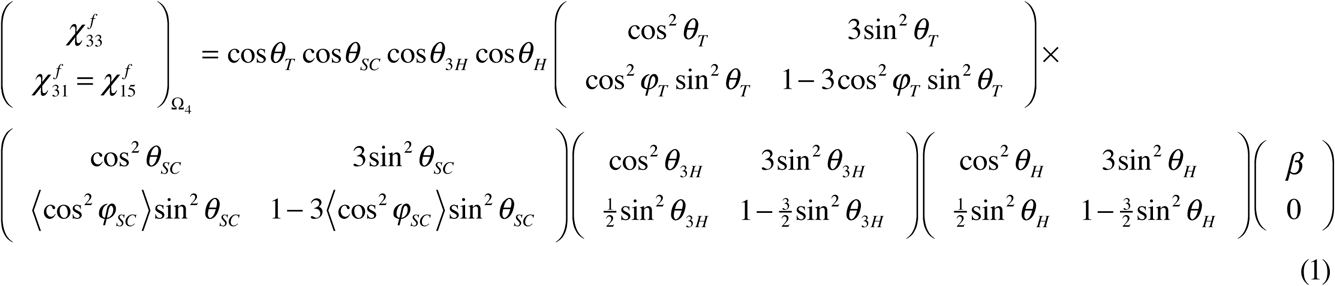

when written in microscope stage coordinate system Ω4 (x_4_, y_4_, z_4_), (see Fig. 1 f). In this equation,*θ*_*H*_, *θ*_*3H*_, *θ*_*SC*_, *θ*_*T*_ are the polar angles of respectively helix (H), triple helix (3H), supercoil (SC) and tilt (T) processes and *φ* _*SC*_ and *φ* _*T*_ are the azimuthal angles of respectively SC and T processes. In this equation, we assume that fibrillar number density of nonlinear dipoles is equal to unity. Since the pitch P_SC_ = 1 μm (20, 21) of the supercoil helix is greater than the transverse PSF (0.4 μm, see Materials and Methods), < > stands for an average over *φ*_*SC*_ at constant *θ*_*SC*_ obtained on the part of the supercoil fibril that is contained in the PSF.

**Figure 1:**
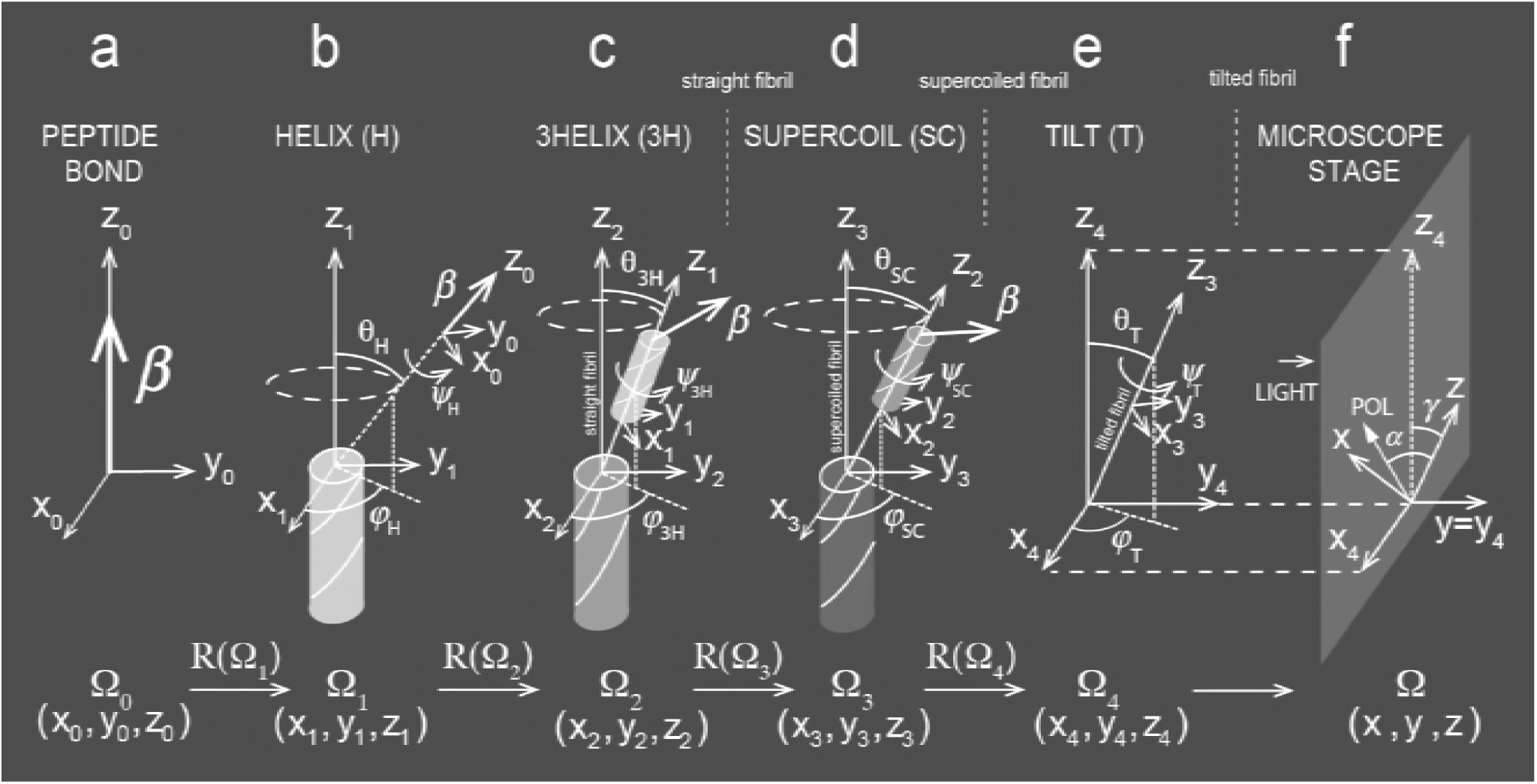
Schematic of hierarchical organization of collagen fibrils. (a) Peptide bond is the dominant contributor to the elementary molecular hyperpolarizability *β* in coordinate system Ω_0_(x_0_, y_0_, z_0_). (b) Cylindrical organization of peptide bond related *β* around main axis z_1_ of singlehelix (H) in coordinate system Ω_1_ (x_1_, y_1_, z_1_). (c) Cylindrical organization of single helix aroundmain axis z_2_ of triple helix (3H) in coordinate system Ω_2_(x_2_, y_2_, z_2_). z_2_ corresponds to the main axis of the straight fibril. (d) Cylindrical organization of triple helix (3H) around main axis z_3_ ofthe supercoiled fibril (SC) in coordinate system Ω_3_(x_3_, y_3_, z_3_). (e) Tilted fibril (T) around direction z_4_ which is in the plane of the microscope stage and which belongs to coordinate system Ω4 (x_4_, y_4_, z_4_). *R*(Ω*i*) is stated for transformation between two successive Ω*i*-1 and Ω*i* coordinate systems (i = 1, 2, 3, 4). The latter is represented in the former with Euler angles (*ψ* _*H*_, *θ*_*H*_, *φ*_*H*_), (*ψ* _3H_, *θ*_3H_, *φ*_3H_), (*ψ* _*SC*_, *θ*_*SC*_, *φ*_*SC*_), (*ψ* _*T*_, *θ*_*T*_, *φ*_*T*_) for respectively i = 1, 2, 3, 4. (f) Microscope stage in fixed laboratory coordinate system Ω(*x, y, z*). γ and *α* are angles between respectively z_4_ direction and input polarization (POL) with z direction. Light propagates in y = y_4_ direction.

Based on staggering and cross-linking properties of fibrillar collagen molecules and the non-centrosymmetric requirement for the development of the SHG process, we assume that each pixel consists of N_f_ identical fibrils, such that theoretical anisotropy parameter of each pixel is 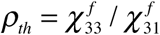 and SHG intensity *I* ^*2ω*^of each pixel is given by the usual formula (1, 5-14, 22-28)

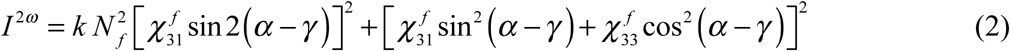

In this equation, *γ* and *α* are angles of respectively z_4_ and incident laser input polarization (POL) directions with fixed z direction of the microscope stage (see Fig. 1 f) and the scaling factor k is proportional to the setup geometry and to the square of the incident IR laser intensity.

In order to take account of a possible fibrillar disorder that may be present in collagen tissues, we next assume that each pixel of the SHG image may contain fibrils with a different fibrillar architecture i.e. different *θ*_*SC*_, *θ*_*T*_ values. Three simple cases that accurately simulate the dispersion of *ρ* values observed in collagen-rich tissues such as tendon, skin and liver vessels are considered. A diagram regrouping the three cases is shown in Fig. 2 where four pixels and three fibrils per pixel (N_f_ = 3) are displayed for both straight and supercoiled fibrils. So we assume that the fibrillar architecture may be due to a constant change (case 1) or to a dispersion (case 2) of tilt angle *θ*_*T*_ from in plane position (*θ*_*T*_ = 0). In these two cases, supercoiled fibrils are regular and characterized by a constant supercoil angle *θ*_*SC*_. For supercoiled fibrils, we hypothesize that in addition to the two previous cases, fibrillar architecture may be due to a variation of the supercoil angle (std *θ*_*SC*_ ≠0) along main fibril axis (case 3), producing irregular supercoiled fibrils with an accordion-like shape. It is worth noting that in cases 2 and 3, values of 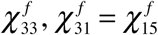 and *ρ* _*th*_ vary with *θ*_*T*_ and *θ*_*SC*_ leading to a greater dispersion of these parameters in case of increased fibrillar disorder.

**Figure 2.**
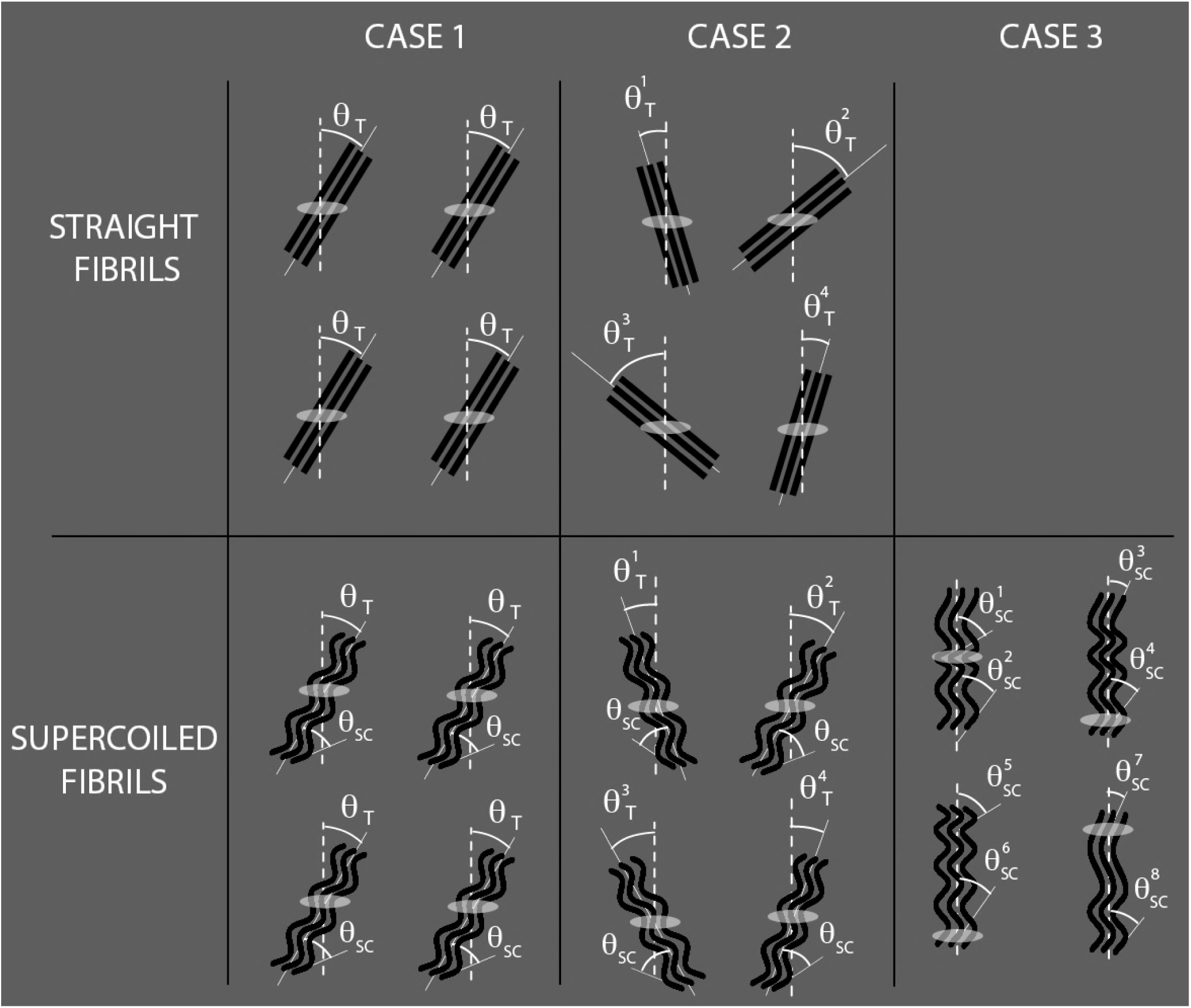
Intra and inter pixel fibrillar architecture. For example, 4 pixels and 3 fibrils per pixel (N_f_ = 3) are considered in each case (1, 2, 3) for straight and supercoiled fibrils. Note that the PSF that materializes the pixel is indicated by the ellipse and that the dash line corresponds to the plane of the microscope stage. Note also that each pixel is located at the center of the PSF. (case 1) Constant tilt angle *θ*_*T*_from in plane position(*θ*_*T*_ = 0). (case 2) variation of tilt angle 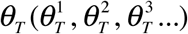 around *θ* _*T*_ = 0. Supercoiled fibrils are regular with constant supercoil angle*θ*_*sc*_ (case 3) Variation of supercoil angle 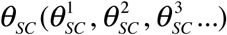 along main fibril axis producing accordion-like supercoiled fibrils.

Since we have recently shown that the experimental anisotropy parameter of each pixel *ρ*_exp_, obtained by fitting P-SHG data with Eq. 2 is highly dependent on Poisson photonic shot noise of the detection system (14), it is of paramount importance to take Poisson noise into account in the calculation of the theoretical anisotropy parameter called *ρ*_*poiss*_ in the following. *ρ*_*poiss*_ *is* obtained from a random P-SHG intensity curve generated from Eq. 2 with Poisson parameter proportional to *I* ^*2ω*^ (see Supporting Material S3 for more details). In summary, three anisotropy parameters *ρ* exp, *ρ* _*th*_ and *ρ* _*poiss*_ are determined for each pixel of the SHG image: 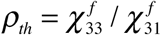 is directly calculated from Eq. 1 while *ρ*_exp_and *ρ*_*poiss*_are obtained by LLS fitting of Eq. 2 on respectively experimental and Poisson noise P-SHG intensity curves (details of the LLS fitting procedure is also described in Supporting Material S3).

## MODEL SIMULATION RESULTS

In order to rigorously explain dispersion of experimental *ρ*_exp_ values in collagen-rich tissues, we model in the following the impact of the inter-pixel fibrillar disorder on the dispersion of theoretical *ρ*_*poiss*_ values for the three cases of fibrillar arrangements described above. We first consider the case of straight fibrils in Fig. 3, then that of the supercoiled fibrils in Fig. 4. The three cases of fibrillar architecture are recalled in inset for each figure. Since we assume that there is no intra-pixel fibrillar disorder, only one fibril is represented for each pixel and therefore, each fibril represented in each inset belongs to a different pixel. Theoretical results can be summarized as follows.

**Figure 3.**
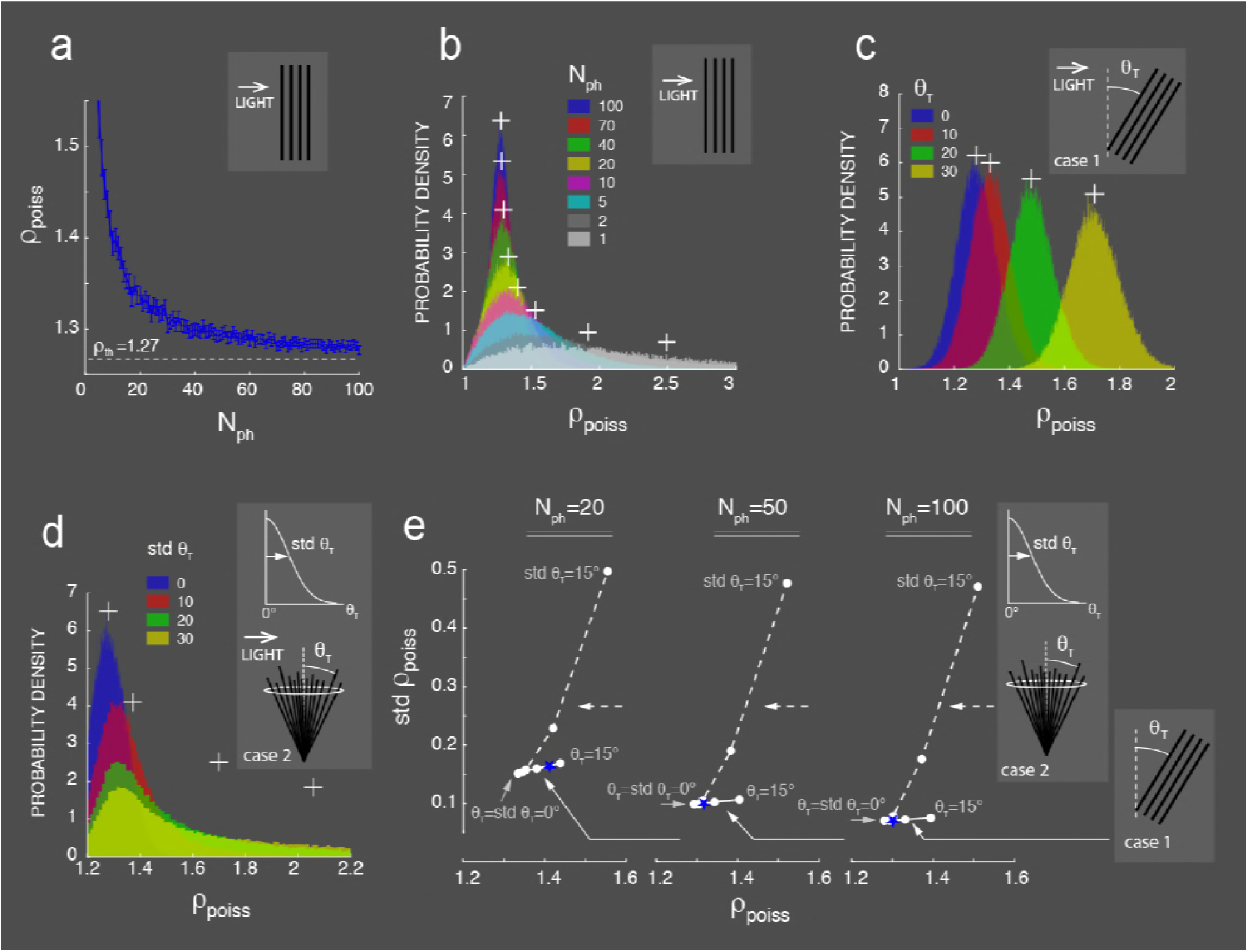
Theoretical calculation of anisotropy parameter *ρ*_*poiss*_ for straight fibrils. (a) Evolution of *ρ*_*poiss*_ for in plane fibrils as a function of the P-SHG stack mean photon number per pixel N_ph_. The curve is obtained with 1000 iterations for each value of N_ph_. Bar graphs represent 97% confidence interval (± 2 SEM) and horizontal dotted line is *ρth* obtained for *θ*_*H*_ = 53° and *θ*_3H_ = 12°. (b-d) Normalized histograms as a stair-case shaped estimation of the probability density function of anisotropy parameter *ρ*_*poiss*_ for (b) in plane ordered fibrils as a function of N_ph_ (c) out of plane fibrils with constant tilt angle *θT* (case 1). Simulation is also obtained for the particular case of fibrils remaining in the plane of incidence of the IR laser beam, *φ*_*T*_ = 90° (d) disordered fibrils with normal distribution of tilt angles *θT* for different values of std *θ*_*T*_. *θT* is chosen randomly for each iteration within a truncated normal distribution to avoid negative values. Azimuthal angles *φ*_*T*_ are chosen randomly (case 2). Note that for each histogram (b-d), mean value of *ρ*_*poiss*_ is indicated at the corresponding abscissa by a white cross positioned at the level of the maximum value of the histogram on the ordinate axis. Each colored histogram corresponds to different values of N_ph_ (b), *θT* (c) and std *θT* (d) as indicated in insets. (e) std *ρ*_*poiss*_as a function (c) of mean value of *ρ*_*poiss*_ for N_ph_ = 20, 50, 100. Continuous and dashed line curves correspond respectively to cases 1 and 2 as shown in insets. Note that for each curve, full white circles correspond to values of either *θT* or std *θ*_*T*_ equal to 0°, 5°, 10°, 15° (from left to right). Coordinates of blue asterisks are experimental mean *ρ*_exp_ and std *ρ*_exp_ values obtained for collagen tendon of Fig. 5. Histograms and curves (b-d) are obtained for 105 iterations and histograms (c, d) are obtained for N_ph_ = 100.

**Figure 4.**
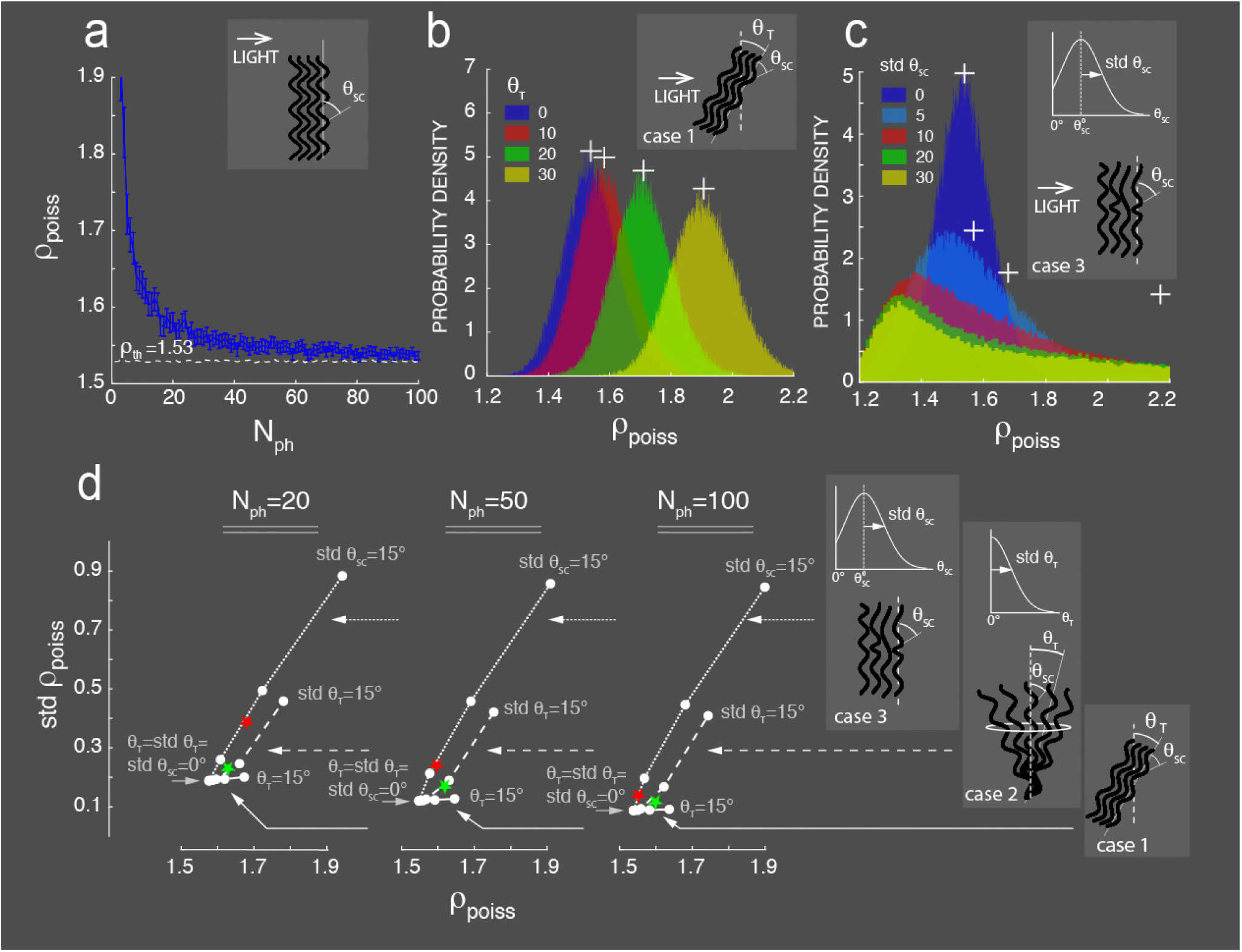
Theoretical calculation of anisotropy parameter *ρ*_*poiss*_ for supercoiled fibrils. (a) Evolution of *ρ*_*poiss*_ as a function of N_ph_ for regular supercoiled fibrils *θ*_*SC*_ = 17°. The curve is obtained over 1000 iterations for each N_ph_. Bar graphs represent 97% confidence interval (± 2 SEM). Horizontal dotted line is *ρ*_*th*_ = 1.53. (b, c) Normalized histograms (as a stair-case shaped estimation of the probability density function) of anisotropy parameter *ρ*_*poiss*_for (b) out of plane regular supercoiled fibrils as a function of constant tilt angle *θT* (case 1). Simulation is also obtained for the particular case of fibrils remaining in the plane of incidence of the IR laser beam, *φ*_*T*_ = 90° (c) accordion-like disordered supercoiled fibril with normal distribution of *θ*_*SC*_ around *θ*_*SC*_ = 17° and different values of std *θ*_*SC*_ (case 3). The simulation is obtained with an average over *φSC* (not shown) obtained on the part of the supercoiled fibril contained in the PSF (see Supporting Material S2 for more details). Note that the normal distribution is truncated to avoid negative values of *θ*_*SC*_. For each histogram in panels b and c, mean value of *ρ*_*poiss*_ is indicated at the corresponding abscissa by a white cross-positioned at the level of the maximum value of the distribution on the ordinate axis. Note that mean value of *ρ*_*poiss*_ for std *θ*_*SC*_= 30° in panel c is out of range. (d) std *ρ*_*poiss*_ as a function of mean value of *ρ*_*poiss*_ for N_ph_ = 20, 50, 100. Continuous, dashed and dotted line curves correspond respectively to cases 1, 2 and 3. Note that for each value of N_ph_, full white circles correspond to values of either *θT*, std *θT* or std *θSC* equal to 0°, 5°, 10° and 15° (from left to right). Coordinates of green and red asterisks are experimental mean *ρ*_exp_ and std *ρ*_exp_ values obtained for respectively skin and control mouse liver vessel of Fig. 6.Histograms and curves (b-d) are obtained for 105 iterations. Histograms (b, c) are obtained for N_ph_ = 100.

For straight fibrils, the case of pixels with in plane ordered fibrils (*θ*_*T*_ = 0) is considered first. Tendon collagen could be the closest tissue corresponding to this case. The evolution of anisotropy parameter *ρ*_*poiss*_ as a function of the P-SHG stack mean photon number per pixel N_ph_ is shown in Fig. 3 a. Simulation is obtained with *θ*_*H*_ = 53°, *θ*_*3H*_ = 12° corresponding to the experimental mean values found in rat tendon for N_ph_ = 100 (see Table 1). The curve decreases quasi exponentially with increase N_ph_ toward *ρ* _*th*_ = 1.27 and a bias due to Poisson noise is clearly visible below N_ph_ = 40. Normalized histograms highlight the decrease of both mean (white crosses) and dispersion of *ρ*_*poiss*_ with increasing N_ph_ (Fig. 3 b). Note that the effect of *θ* _*H*_ and *θ*_*3H*_ on *ρ*_*poiss*_ is not shown in Fig. 3 a and b. Increasing their values within the range of Table 1 results in respectively decrease and increase of *ρ*_*poiss*_. More importantly, we noticed that their dispersion within the same range had a minor impact on the dispersion of *ρ*_*poiss*_ for both straight and supercoiled fibrils and for this reason *θ*_*H*_ and *θ*_*3H*_ are considered constant in the following. The case of pixels with out of plane straight fibrils and constant tilt angle *θ*_*T*_ ≠ 0 (case 1) is shown in Fig. 3 c. Histograms of *ρ*_*poiss*_are characterized by an increase of both mean (white crosses) and dispersion of *ρ*_*poiss*_ with increasing *θ*_*T*_ from 0° to 30°.

**Table 1:**
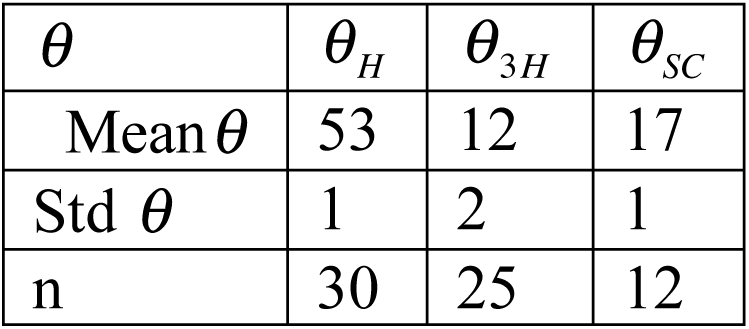
Statistical hierarchical angles in degrees for rat tendon and skin obtained from n samples at Nph =100.

The case of pixels with disordered straight fibrils with random tilt angles std *θ*_*T*_ ≠0 (case 2)around *θ*_*T*_ = 0is shown in Fig. 3 d. Histograms of *ρ*_*poiss*_ are characterized by a larger increase of both mean (white crosses) and dispersion of *ρ*_*poiss*_ with increasing std *θ*_*T*_ from 0° to 30°. Simulation is obtained for angles *θ*_*T*_. that are distributed within a normal distribution around *θ*_*T*_ = 0° as indicated in the inset

Correlation between *ρ*_*poiss*_and std *ρ*_*poiss*_is more precisely shown in Fig. 3 e for different values of N_ph_ (20, 50, 100) and for the two cases of fibrillar architecture considered: constant tilt angle *θ*_*T*_ ≠ 0 (continuous line curves, case 1) and random tilt angle, std *θ*_*T*_≠0 (dashed line curves, case 2) and for values of either *θT* or std *θ*_*T*_ varying from 0° to 15°.

The major finding of the simulation is that dispersion of *ρ*_*poiss*_(std *ρ*_*poiss*_) increases more rapidly for fibrils with random tilt angles (case 2) compared to fibrils with constant ones (case 1) as clearly shown by the difference of slope between the continuous and dashed line curves.

For supercoiled fibrils, the simple case of in plane ordered fibrils (*θ*_*T*_ = 0) with regular supercoil angle *θ*_*SC*_ = 17° is considered first. Skin collagen could be the closest tissue corresponding to this case. The evolution of *ρ* _*poiss*_ as a function of N_ph_ is shown in Fig. 4 a. The simulation is obtained with *θH* = 53°, *θ*_3H_ = 12° and *θ*_*SC*_ = 17° corresponding to mean values found in rat skin for N_ph_ =100 (see Table 1). As in tendon, the curve decreases with increasing N_ph_ toward higher *ρth* = 1.53. Similarly, a bias due to Poisson noise is also observed below N_ph_ = 40. The case of out of plane regular supercoiled fibrils with constant tilt angle *θ*_*T*_ ≠ 0 (case 1) is shown in Fig. 4 b. Tilting the fibrils has a similar increasing effect on *ρ*_*poiss*_ and std *ρ* _*poiss*_ as in tendon (compare with Fig. 3 c).

The case of supercoiled fibrils with fibrillar disorder due to dispersion of tilt angles std *θ*_*T*_ ≠ 0 (case 2) is represented later in Fig. 4 d. The histograms are similar to that obtained for straight fibrils in Fig. 3 d except that they are shifted to higher values.

Finally, the case of fibrillar disorder due to variation of supercoil angles std *θ*_*SC*_ ≠ 0 around mean position *θ*_*SC*_ = 17° (case 3), producing irregular supercoiled fibrils with an accordion-like shape is considered next in Fig. 4 c. Histograms of *ρ*_*poiss*_are characterized by a more important increase of both mean (white crosses) and dispersion of *ρ* _*poiss*_ as compared to the two previous cases.

Correlation between *ρ*_*poiss*_and std *ρ*_*poiss*_is more precisely shown in Fig. 4 d for different values of N_ph_ (20, 50, 100) and for the three cases of fibrillar architecture considered: constant tilt angle *θ*_*T*_ ≠ 0 (continuous line curves, case 1); random tilt angle, std *θ*_*T*_ ≠ 0 (dashed line curves, case 2) and random supercoil angle, std *θ*_*SC*_ ≠ 0 (dotted line curves, case 3) and for values of either *θ*_*T*_, std *θ*_*T*_ or std *θ*_*SC*_ varying from 0° to 15°. Compared to straight fibrils (see Fig. 3 e) continuous and dashed line curves are shifted while dotted ones are steeper.

Altogether, these results show that (i) for fibrils with constant tilt angle *θ*_*T*_ ≠ 0 (case 1), mean value of *ρ*_*poiss*_significantly increases from straight to supercoiled fibrils while dispersion of *ρ* _*SC*_ is dominated by Poisson shot noise (std *ρ*_*poiss*_~ 0.1 at N _*ph*_ = 100) (ii) for disordered straight and supercoiled fibrils std *θ*_*T*_ ≠ 0(case 2) and std *θ*_*SC*_ ≠ 0 (case 3), both mean and dispersion of *ρ* _*poiss*_ increase with increasing dispersion of *θ*_*T*_ and *θ*_*SC*_ but the effect of the latter is more drastic.

## EXPERIMENTAL RESULTS

In order to determine the origin of the dispersion of experimental anisotropy parameters *ρ*_exp_, we choose to correlate our architecture-based model of theoretical anisotropy parameters *ρ* _*poiss*_ to in situ experimental ones *ρ*_exp_ obtained from typical examples of respectively straight (*θ* _*sc*_ = 0,tendon) and supercoiled(*θ*_*SC*_ ≠ 0,skin, liver vessels) collagen fibrils. All following theoretical simulations were obtained using hierarchical angles *θ*_*H*_ = 53° and *θ*_*3H*_ = 12° corresponding to the best match between *ρ* _*poiss*_ and *ρ* exp found in rat tendon and skin for N_ph_ = 100 (see Table 1).

For rat extensor digitorum longus tendon, P-SHG image and map of *ρ*_exp_ are shown respectively in Fig. 5 a and b. Distribution of *ρ*_exp_ decreases with N_ph_ (Fig. 5 c) as expected from theoretical analysis (see Fig. 3) For this tissue, we obtain *ρ*_exp_ = 1.33 ± 0.11 over the entire SHG image (see also Table 2). Mean and dispersion of *ρ*_exp_, *ρ*_*poiss*_, *ρ*_*th*_ are shown in Fig. 5 d and e as a function of N_ph_. Theoretical simulations of *ρ* _*poiss*_ (blue curves) and *ρ* _*th*_ (white curves) are obtained considering in plane ordered straight fibrils (*θ*_*T*_ = 0). While mean values are slightly different in particular for N_ph_ < 80 (Fig. 5 d, red and blue curves), std *ρ* exp and std *ρ* _*poiss*_ are very similar (Fig. 5 e, red and blue curves) suggesting that dispersion of *ρ*_exp_ is mainly due to Poisson noise considering in plane ordered straight fibrils. However, the mismatch of histograms of *ρ*_exp_ and *ρ*_*poiss*_(Fig. 5 f, red and blue histograms) indicates that some fibrils are probably tilted and therefore out of plane (*θ*_*T*_ ≠ 0), corresponding to case 1 of inter pixel fibrillar architecture (see Fig. 2). As the dispersion of the experimental *ρ* exp values decreases with N_ph_ (see Fig. 5 c and e), the fibrillar architecture is estimated for each value of N_ph_, i.e. between pixels having an identical value of N_ph_. The mismatch between distributions of *ρ*_exp_and *ρ*_*poiss*_is calculated for each interval of ten photons by minimizing the Kolmogorov-Smirnov distance *dk* (29) between the two empirical cumulative distribution functions associated to *ρ*_exp_ and *ρ*_*poiss*_as a function of the tilt angle *θ*_*T*_ (Fig. 5 g, green curve). From dk, we have defined a correlation coefficient *R*_*k*_ = 1 - *d*_*k*_ (*R*_*k*_ ∈[0 1]) that is used to determine the best match between distributions of *ρ*_exp_ (red curves) and *ρ*_*poiss*_(magenta curves) as shown in fig. 5 g-i. As a result of this correction, we obtain *θ*_*T*_ ~ 7.3° over the entire SHG image and R_k_ increases from 0.89 to 0.99 without and with correction of tilt angle *θ* _*T*_. Altogether, these results show that rat tendon is characterized by preponderant ordered straight fibrils and that dispersion of *ρ*_exp_ values is mainly described by effect of Poisson noise. Small contribution of fibrillar arrangement associated with constant tilt angles (case 1) is often observed for pixels with low N_ph_ values.

**Figure 5.**
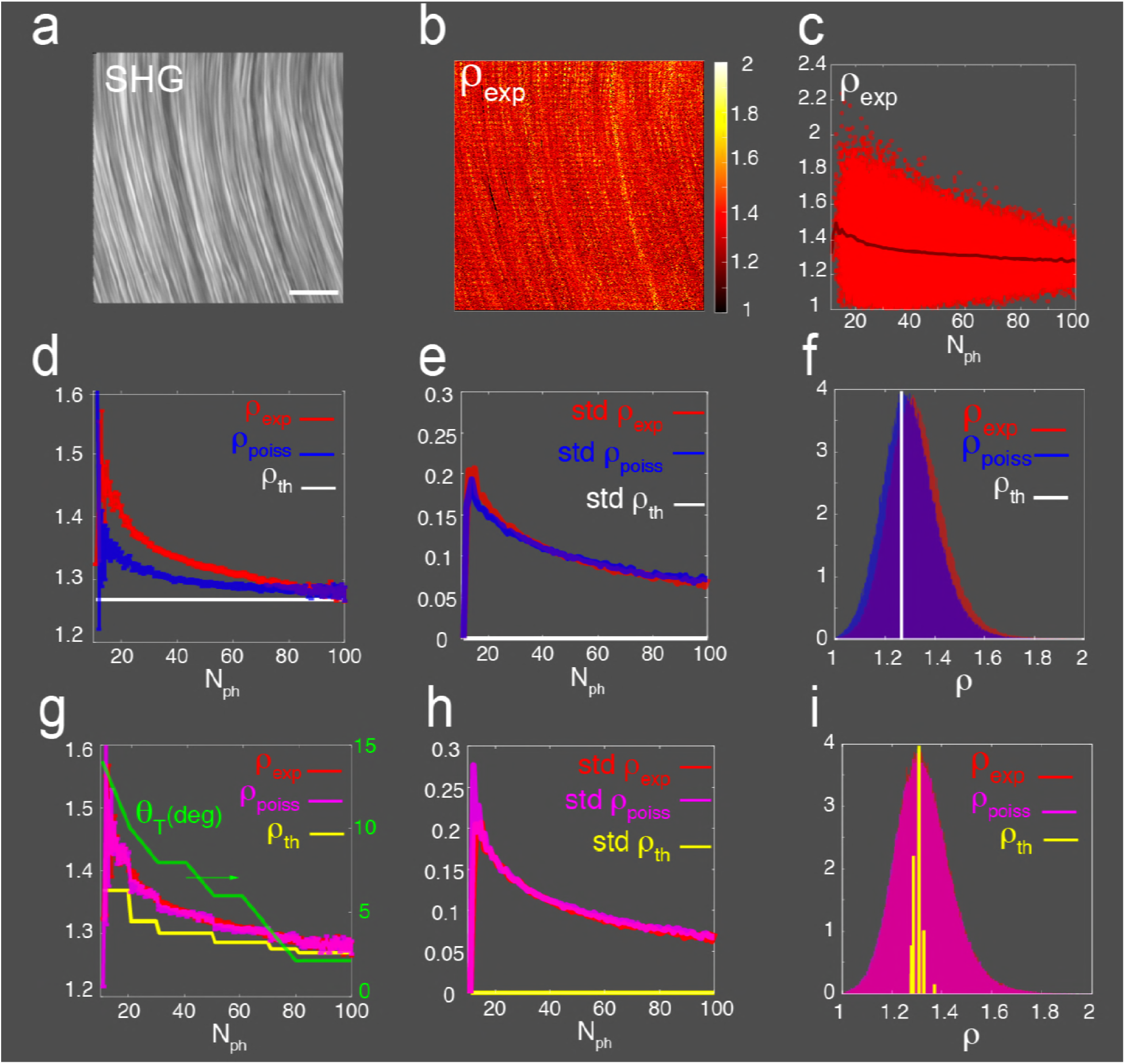
P-SHG image analysis of rat EDL tendon. (a) Typical 512×512 pixels SHG image obtained from average of the P-SHG stack. Scale bar is 10 μm. (b) Map of experimental anisotropy parameter *ρ*_exp_ for each pixel of the SHG image. (c) Distribution of *ρ*_exp_ as a function of N_ph_. Full line corresponds to mean values of *ρ*_exp_. (d) Mean ± 2 SEM curves of *ρ*_exp_ (red line curve), theoretical without *ρ* _*th*_ (white line curve) and with Poisson noise *ρ*_*poiss*_ (blue line curve) as a function of N_ph_. Bar graphs represent 97% confidence interval (± 2 SEM). (e) std *ρ*_exp_, std *ρ* _*th*_, std *ρ*_*poiss*_ curves as a function of N_ph_. (f) Normalized histograms of *ρ*_exp_, *ρ* _*th*_, *ρ*_*poiss*_ distributions. Theoretical curves of d, e, f in white and blue color are obtained considering in plane ordered straight fibrils *θ*_*T*_ = 0°. Kolmogorov-Smirnov correlation coefficient between *ρ*_exp_ and *ρ*_*poiss*_ distributions is R_k_ = 0.89. (g, h, i) have the same meaning as in (d, e, f) but theoretical curves in yellow and magenta color are obtained for out of plane fibrils *θ*_*T*_ ≠ 0°. *θ* _*T*_ was calculated by minimizing the Kolmogorov-Smirnov distance d_k_ between the two empirical cumulative distribution functions associated to *ρ*_exp_ and *ρ*_*poiss*_ for each interval of ten photons (green curve of panel g with values on right y-axis). The overall correction corresponds to a Kolmogorov-Smirnov mean correlation coefficient *R*_*k*_ = 1 - *d*_*k*_ = 0.99 between *ρ*_exp_ and *ρ*_*poiss*_ distributions associated with a mean tilt angle *θ*_*T*_ = 7.3° over the entire SHG image (see also Table 2). Note the good superposition of the red and magenta curves as well as red and magenta histograms.

**Table 2:** Statistical parameters for different collagen-rich tissues: rat tendon, rat skin, rat media, control (Oil-W8) and fibrotic (CCl4-W8) mouse liver vessels. All values are mean values averaged over number of samples n.

|  | Rat Tendon | Rat Skin | Rat Media | Oil-W8 | CCl<sub>4</sub>-W8 |
| --- | --- | --- | --- | --- | --- |
| $\rho_{\text{exp}}$ | 1.35 | 1.63 | 1.67 | 1.68 | 1.67 |
| Std $\rho_{\text{exp}}$ | 0.14 | 0.19 | 0.30 | 0.38 | 0.36 |
| N<sub>ph</sub> | 47 | 48 | 22 | 23 | 25 |
| $\theta_T$ | 7 | ~ 0 | ~ 0 | ~ 0 | ~ 0 |
| Std $\theta_T$ | ~ 0 | 9.6 | ~ 0 | ~ 0 | ~ 0 |
| $\theta_{SC}$ | $\triangleq 0$ | 17 | 17 | 17 | 17 |
| Std $\theta_{SC}$ | $\triangleq 0$ | ~ 0 | 4.5 | 6.3 | 5.4 |
| R<sub>k</sub> | 0.99 | 0.99 | 0.98 | 0.97 | 0.96 |
| n | 18 | 9 | 7 | 24 | 23 |

We next extend our study to tissues characterized by supercoiled collagen fibrils such as skin and liver vessels. In a previous study, we showed that the distribution of mean values of experimental anisotropy parameters *ρ* exp was well described by our theoretical model, but we did not explain their dispersions rigorously (15). Despite taking into account both Poisson noise and fibrillar disorder due to the tilt of fibrils (std *θ*_*T*_ ≠ 0) corresponding to case 2 of inter pixel fibrillar architecture (see Fig. 2), dispersion of experimental *ρ* exp values were always greater than theoretical ones *ρ* _*poiss*_ (15). Based on our new theoretical modeling indicating that dispersion of *ρ*_*poiss*_ can be further enhanced by introducing the dispersion on supercoil angles (std *θ*_*SC*_ ≠ 0) corresponding to case 3 of inter pixel fibrillar architecture (see Fig. 2), we next test this hypothesis.

For rat skin, P-SHG image and normalized histogram of *ρ* exp (red color) are shown respectively in Fig. 6 a and b. Histogram is characterized by *ρ* exp = 1.62 ± 0.17 with mean N_ph_ ~ 50 over the entire SHG image (see also Table 2). Mean *ρ* exp value is significantly (p<0.001) above that in tendon. However, std *ρ* exp = 0.17 is very close to std *ρ* _*poiss*_ = 0.13 obtained when considering in plane and ordered regular supercoiled fibrils *θ*_*T*_ = 0°, *θ*_*SC*_ = 17°. Extrapolating from the experimental values (green asterisks), the theoretical curves of Fid. 4 d suggest that dispersion of *ρ* exp should originate from dispersion of tilt angles *θ*_*T*_ (case 2) rather than supercoil angles *θ*_*SC*_ (case 3). Theoretical simulation reveals that the best match (R_k_ = 0.99) between *ρ*_exp_ (red color) and *ρ*_*poiss*_(magenta color) histograms (Fig. 6 b) is obtained for a mean dispersion of tilt angles std *θ*_*T*_ ~ 13° calculated over the entire SHG image and corresponding to *ρ* _*th*_ (yellow color) histogram (see also Fig. S3 of Supporting Material S4 for more details). These results suggest that the dispersion of *ρ* expvalues in rat skin is mainly due to both Poisson noise with a contribution due to fibrillar disorder associated with a dispersion of tilt angles (std *θ*_*T*_ ≠ 0, case 2).

**Figure 6.**
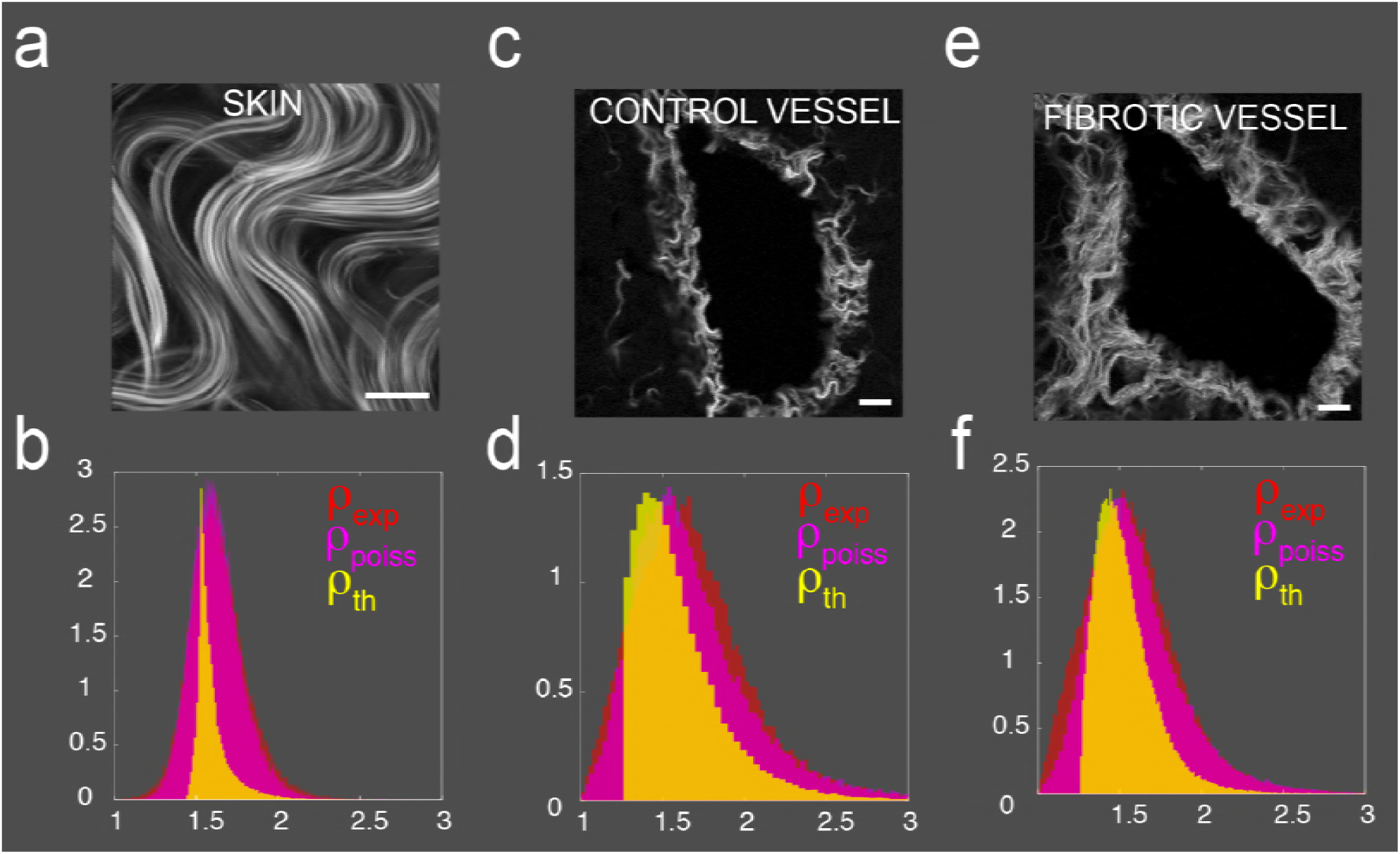
P-SHG image analysis of rat skin and mouse liver vessels. (a, b) rat skin. (c, d) control mouse liver vessel. (e, f) CCl_4_-treated mouse liver vessel. (a, c, e) Typical 512×512 pixels SHG images. (b, d, f) normalized histograms for respectively rat skin, mouse control and CCl_4_-treated liver vessels of experimental *ρ*_exp_ (red color), theoretical *ρ* _*th*_ (yellow color) and *ρ*_*poiss*_ (magenta color) distributions. *ρ* _*th*_ (yellow color) gives the best correspondence with the greatest Kolmogorov-Smirnov correlation coefficient R_k_ between *ρ*_exp_ (red color) and *ρ*_*poiss*_(magenta color) distributions. (b) R_k_ = 0.99 between *ρ*_exp_ = 1.62 ± 0.17 and *ρ*_*poiss*_= 1.62 ± 0.17 for *θT* = 0° ± 13°, *θ* _*SC*_ = 17° ± 0°. (d) R_k_ = 0.95 between *ρ*_exp_ = 1.70 ± 0.42 and *ρ*_*poiss*_= 1.69 ± 0.41 for *θT* = 0°, *θ* _*SC*_ = 17° ± 7°. (f) R_k_ = 0.95 between *ρ*_exp_ = 1.58 ± 0.30 and *ρ*_*poiss*_= 1.60 ± 0.29 for *θ*_*T*_ = 0°, *θ*_*SC*_ = 16° ± 5°. Scale bars are 10 μm and 20 μm for respectively (a) and (c, e). Moredetails of the fitting procedure are available in Supporting Material S3.

For control mouse liver vessels P-SHG image analysis is shown in Fig. 6 c and d. Normalized histogram of *ρ* exp (red color) is characterized by *ρ*_exp_ = 1.70 ± 0.42 with mean N_ph_ ≠ 23 over the entire SHG image (see also Table 2). Compared to skin, while mean *ρ*_exp_ value is similar (p = 0.1), dispersion is significantly greater (p<0.001) suggesting greater fibrillar disorder. Extrapolating from the experimental values (red asterisks), the theoretical curves of Fid. 4 d suggest that dispersion of*ρ* exp should originate from dispersion of supercoil angles (std *θ*_*SC*_ ≠ 0, case 3) rather than tilt angles (std *θ*_*T*_ ≠ 0, case 2). Thus, the only way to achieve the best match between dispersion of *ρ* exp and *ρ* _*poiss*_ is to take into account additional disorder of supercoil *θ*_*SC*_ angles assuming accordion-like supercoiled fibrils. Theoretical simulation reveals that the best match (R_k_ = 0.95) between *ρ*_exp_ (red color) and *ρ*_*poiss*_ (magenta color) histograms (Fig. 6 d) is obtained for *θ*_*SC*_ = 17.0° ± 7° calculated over the entire SHG image and corresponding to *ρ*_*th*_(yellow color) histogram (see also Fig. S4 of Supporting Material S4 for more details). Similar disorder of supercoil *θ*_*SC*_ angles is also found in rat aorta media (see Table 2).

We have previously shown that fibrosis in mouse liver vessels is characterized by a significant reduction in the mean value of *ρ*_exp_ when taking into account pixels with low N_ph_ (5<N_ph_<50), compared to that obtained in mouse liver control and this has been interpreted as a reduction of fibrillar disorder (15). However, this interpretation did not consider the difference between the dispersions of *ρ*_exp_ and *ρ*_*poiss*_, that we address in the following. Fibrotic CCl_4_-treated mouse liver vessels P-SHG image analysis is shown in Fig. 6 e and f. Normalized histogram of *ρ*_exp_ (red color) is characterized by *ρ*_exp_ = 1.58 ± 0.30 with mean N_ph_ ≠ 25 over the entire SHG image (see also Table 2). While mean values between control and fibrotic vessels are not significantly different (p = 0.45), dispersion of *ρ*_exp_ is significantly reduced (p<0.05) in fibrotic vessel compared to control one. Theoretical simulations reveal that the reduction of the dispersion of *ρ*_exp_ in fibrotic vessel is due to a significant (p<0.001) reduction of the inter pixel disorder of the supercoil angles *θ_SC_* from 7° to 5° (see also Table 2), that corresponds to the best match (R_k_ = 0.95) between *ρ*_exp_ (red color) and *ρ*_*poiss*_ (magenta color) histograms of Fig. 6 f (see also Fig. S4 of Supporting Material S4 for more details).

Altogether, these results suggest for the first time to our knowledge that distribution of experimental anisotropy parameter *ρ*_exp_ in vessels is mainly driven by a variation of supercoil angles within a single accordion-like supercoiled fibril. This disorder is reduced in fibrotic livers, suggesting more efficient cross-linking and an increase stiffening compared to control vessels.

## DISCUSSION

Our analytical calculation of the second-order nonlinear optical susceptibility tensor *χ* ^(2)^ in collagen-rich tissues is based on the approximation of a single element *β*_33_ = *β*in the associated molecular hyperpolarizability tensor *β* ^(2)^. Within this approximation, we found that the best estimated value of single helix angle (H) determined from collagen tendon is *θ*_*H*_ = 53 ± 1° (see Table 1). This value is similar to that previously found in collagen tendon (5, 30, 31) but is about 7° higher than value deduced from X-rays diffraction studies (20, 22). Although peptide bonds have a major contribution to SHG signal (5, 6, 11, 26), this discrepancy suggests other contributions to SHG signal as previously reported (9, 13, 32). Interestingly, our assumption of an equivalent unique single molecular hyperpolarizability tensor element *β* is further validated by experimental results showing that *χ* _31_ = *χ* _15_ in several collagen tissues (see Supporting Material S1). Moreover, our estimation of triple helix (3H) angle *θ* _*3H*_ = 12 ± 2°in collagen tendon is in agreement with X-rays diffraction studies (20). We also assumed that supercoiled fibrils characteristics of skin and vessels, correspond to an additional helical supercoil (SC) arrangement obtained from straight fibrils with a mean supercoil angle *θ*_*SC*_ = 17°. This was necessary to accurately simulates distribution of experimental anisotropy parameter *ρ*_exp_ in these tissues as previously reported (12). In this context, the two independent nonlinear optical susceptibility coefficients *χ*_33_, *χ*_31_ contributing to the SHG signal as well as the associated anisotropy parameter *ρ* _*th*_ = *χ*33 / *χ*31 were calculated as a function of the known hierarchical poly-helical assembly of the collagen straight or supercoiled fibrils. In order to correlate dispersions of experimental and theoretical distributions of anisotropy parameters *ρ*_exp_ and *ρ*_*poiss*_, Poisson photonic noise and fibrillar architecture have been considered. Thus, by minimizing the Kolmogorov-Smirnov distance between the two empirical cumulative distribution functions associated to *ρ*_exp_ and *ρ*_*poiss*_, fine fibrillar arrangement has been deduced in different physio-pathological collagen tissues (tendon, skin, liver vessels). We considered only fibrillar disorder originating from either tilt (case 2) or supercoil angular (case 3) dispersions resulting from pixel to pixel random variation. Since we observed that the discrepancy between distributions of *ρ*_exp_ and *ρ*_*poiss*_decreases with number of photons N_ph_ in all tissues (see Figs 5, S3, S4, S5 d and e), we assumed that the fibrillar disorder also decreases with N_ph_. We therefore fitted the experimental data for each N_ph_ value with a Kolmogorov-Smirnov test procedure to determine fibrillar disorder for each pixel of a 512×512 SHG image. We found that, for ordered straight and regular supercoil fibrils as tendon and skin, dispersion of *ρ*_exp_ is mainly due to Poisson shot noise. However, we found that the dispersion of *ρ*_exp_ in skin is slightly larger and associated to a slight dispersion of the tilt angles std *θ*_*T*_ ≠ 10°(see Table 2).

The novelty of this study is that dispersion of *ρ*_exp_ values in liver vessels and aorta media might originate from variation of supercoil *θ* _*SC*_ angles along fibrillar main axis. Regular supercoiled fibrils with mean values of 15° ≤ *θ*_*SC*_ ≤ 22° have been observed by ultrastructural SEM studies in skin cornea and vessels, however dispersion of *θ* _*SC*_ has not been reported to our knowledge (19). While P-SHG studies have also reported the presence of regular supercoiled fibrils with constant supercoil angle *θ*_*SC*_ in the same tissues (12, 15), our present study suggests that accordion-like supercoiled fibrils characterized by dispersion of supercoil angles *θ* _*SC*_ are responsible of the dispersion of *ρ*_exp_ values in liver vessels. This assumption is supported by the fact that experimental values for control mouse live vessels (red asterisks) are on the dotted line curves of Fig. 4 d for all values of N_ph_. For this tissue, we obtain std *θ*_*SC*_ = 6.3° calculated over the entire SHG image. Interestingly, similar results are found in rat aorta media std *θ*_*SC*_ = 4.5° (see Table 2). For CCl_4_-treated mouse liver vessels, results show a significant (p<0.05) reduction of the dispersion of the supercoil angles, std *θ*_*SC*_ = 5.4°, compared to control ones (see Table 2). This result is in agreement with our previous results suggesting less disorder in CCl_4_-treated fibrotic vessels (15). While in that study we focused our analysis on the variations of the mean values of *ρ*_exp_, in the present work, we accurately consider their dispersions. It turns out that dispersion of supercoil angles *θ* _*SC*_ well described both mean and standard deviation of *ρ*_exp_ values in liver vessels. Altogether, these results suggest that liver fibrosis is characterized by a remodeling of collagen fibrils favoring increase fibrillar alignment. Since it has been shown that collagen fibrillar alignment is involved in cross-linking and stiffening of the extracellular matrix (33-36), this remodeling will result in an increase in rigidity of vascular wall and a subsequent rise in intra-portal tracts and central veins tone as already suggested (37).

We also noticed that for most collagen tissues studied, the average number of photons N_ph_ for a SHG image is between 20 and 50 (see Table 2) and that the lowest photon numbers are associated with the more disordered tissues. In this range of photon numbers, the Poisson noise impacts the values of the experimental anisotropy parameter *ρ*_exp_ and this concerns both the dispersion and the bias of its values (see Figs 3 a and 4 a). However, thresholding above 50 photons to avoid Poisson noise will result in a great loss of information regarding the estimate of the fibrillar disorder. On the other hand, increasing the acquisition time to improve the signal-to-noise ratio makes it possible to reduce the dispersion of *ρ*_exp_ values, but the disadvantage is probably the presence of more artifacts related to the mechanical drift. For example, with our experimental setup, the acquisition of a SHG image with a duration of one second gives 20 photons per pixel due to the 5MHZ bandwidth limit of the high sensitivity single photon GasAsP photomultiplier (H7421-40) provided by Hamamatsu. In a usual P-SHG acquisition, a compromise needs to be done between acquisition duration and signal to noise ratio. We found that a maximum photons number around 100 is a good compromise. In any case, even at low photon numbers, there is no difficulty in distinguishing between the dispersion of *ρ*_exp_ values between straight and supercoiled fibrils (see Figs 3 a and 4 a). Overall, our study indicates that an accurate estimate of fibrillar disorder from a P-SHG experiment is possible regardless of the number of photons per pixel as long as Poisson noise is considered.

## CONCLUSION

In this study, we have calculated theoretical second-order non-linear optical anisotropy parameter *ρ* = *χ*_33_ / *χ*_31_ for collagen-rich tissues considering the fibrillar disorder. The major contribution of this work, combining theoretical modeling and P-SHG experiment, concerns the structural origin of the dispersion of the values of anisotropy parameter *ρ* in collagen tendon, skin and pathophysiological liver vessels. Our study reveals that fibrillar disorder significantly increases the dispersion of *ρ* values and that its rigorous determination from a P-SHG experiment should consider the Poisson noise of the detection system. The novelty of this study is that dispersion of *ρ*_exp_ values in liver vessels and aorta media might originate from variation of supercoil *θ*_*SC*_ angles along fibrillar main axis suggesting the presence of accordion-like fibrils. In addition, this study paves the way for future modeling correlating dispersion of tissues. *ρ* values, fibrillar disorder and mechanical stiffness of diseases

## SUPPORTING MATERIAL

Supplemental Material S1-S4 are available at http…

## AUTHOR CONTRIBUTIONS

E.S. built the set-up and performed the measurements. F.E. conceived, designed and prepared the biological samples samples. D.R. and J.J.B. developed the theoretical model. D.R. and F.T. designed, performed the experiments, analyzed the data and wrote the manuscript.

## FUNDING AND ACKNOWLEDGMENTS

This work was supported by Région Bretagne, Rennes Métropole, Conseil Général d’Ille-et-Vilaine, Valorial Rennes, Ministère de l’Enseignement Supérieur et de la Recherche and the European Union Federal Funds FEDER.

